# Parallel adaptation to lower altitudes is associated with enhanced plasticity in *Heliosperma pusillum* (Caryophyllaceae)

**DOI:** 10.1101/2022.05.28.493825

**Authors:** Aglaia Szukala, Clara Bertel, Božo Frajman, Peter Schönswetter, Ovidiu Paun

## Abstract

High levels of phenotypic plasticity are thought to be inherently costly in stable or extreme environments, but enhanced plasticity may evolve as a response to novel environments and foster adaptation. *Heliosperma pusillum* forms pubescent montane and glabrous alpine ecotypes that diverged recurrently and polytopically (*parallel evolution*). The specific montane and alpine localities are characterized by distinct temperature conditions, available moisture and light. To disentangle the relative contribution of constitutive versus plastic gene expression to altitudinal divergence, we analyze the transcriptomic profiles of two parallely evolved ecotype pairs, grown in reciprocal transplantations at native altitudinal sites. In both ecotype pairs, only a minor proportion of genes appear constitutively differentially expressed between the ecotypes regardless of the growing environment. Both derived, montane populations bear comparatively higher plasticity of gene expression than the alpine populations that can be considered in this system as ‘ancestor-proxies’. Genes that change expression plastically and constitutively underlie similar ecologically relevant pathways, related to response to drought and trichome formation. Other relevant processes, such as photosynthesis, seem to rely mainly on plastic changes. The enhanced plasticity consistently observed in the montane ecotype likely evolved as a response to the newly colonized niche. Our findings confirm that directional changes in gene expression plasticity can shape initial stages of phenotypic evolution, likely fostering adaptation to novel environments.

**Significance Statement:** Understanding the importance of phenotypic plasticity for fast adaptation to stress is very timely for breeding and current environmental challenges. Our study of an alpine plant in the carnation family evidences an increased level of expression plasticity in early stages of adaptation to hotter and drier habitats.

## Introduction

How phenotypic divergence between conspecific populations arises, possibly leading to local adaptation, stable differentiation and ultimately speciation, is a central question in evolutionary biology. A property of organisms, which can determine population differentiation across heterogeneous environments, is plasticity, i.e., the capacity of a genotype to change its phenotype upon exposure to differing environmental conditions (Schlichting & Pigliucci, 1998). It is still poorly investigated if phenotypic plasticity can promote long-term phenotypic change, and what are the mechanisms behind translating short-term environmental responses into long-term evolutionary states (Stearns, 1989; Sommer, 2020; Stotz *et al*., 2021).

Despite being a necessary property to survive in variable environments, especially for sessile organisms such as plants, plasticity was suggested to be inherently evolutionarily costly when the population reached an adaptive optimum (DeWitt *et al*., 1998; Pál & Miklós, 1999). Moreover, some authors advanced the hypothesis that plasticity might reduce the power of natural selection, representing a dead end of evolution (Charlesworth *et al*., 1982). In stark contrast, phenotypic plasticity has been advanced as a primary object of selection by others (Waddington, 1942; Levis & Pfennig, 2016; Stotz *et al*., 2021), who called for a reconsideration of its importance for the evolutionary process. Adaptive benefits of plasticity have been in certain circumstances documented, for example for traits related to biotic responses (Auld & Relyea, 2011) and abiotic stress (Dudley & Schmitt, 1996; Nicotra *et al*., 2015; Stotz *et al*., 2021; Sole-Medina *et al*., 2022). However, plastic components of some traits can also be neutral or even maladaptive (van Kleunen & Fischer, 2005; Arnold *et al*., 2019a). How and under which conditions plasticity supplies the phenotypic variation later refined by selection remains debated (Flatscher *et al*., 2012; Wund, 2012; Levis & Pfennig, 2016; Arnold *et al*., 2019b; Fox *et al*., 2019).

Plasticity has been linked to local adaptation on rare occasions. Corl *et al*. (2018) showed for example how plasticity at genes controlling skin coloration of the common side-blotched lizard facilitated colonization of dark-soil environments, and suggested that genetic changes in the same genes were shaped by natural selection refining a pre-existing plastic phenotype. Similarly, Levis *et al*. (2018) reported that in spadefoot toad tadpoles adaptive novelty can arise from pre-existing plasticity in diet-related morphological and molecular features. However, the study also uncovered diet-induced maladaptive plasticity affecting mouthpart formation, but also expression of a diet-relevant gene. Selection appears to promote enhanced plasticity in certain conditions but not in others, as shown for example in the waxy bluebell *Wahlenbergia ceracea* Lothian (Campanulaceae; Nicotra *et al*., 2015). In this plant, low-elevation populations show enhanced temperature-induced plasticity, which was found to be more often adaptive compared to populations from higher elevations.

Evidence from natural study systems is required to clarify under which conditions plasticity is favored or hindered by evolution. A meta-study by Barley *et al*. (2021) quantified thermal acclimation capacity across 19 species including arthropods, molluscs, and chordates, showing that within species, marginal populations experiencing the highest thermal conditions had decreased plasticity and acclimation capacity. A negative relationship between plasticity and adaptation to heat extremes was also found in laboratory experiments (Kelly *et al*., 2017; Sasaki & Dam, 2021). These results suggest that there is a trade-off for plasticity at ecological extremes (Chevin & Hoffmann, 2017), such that extreme environments appear to favor phenotypic robustness through *canalization* (Waddington, 1942). On the other hand, evolve and resequence studies on *Drosophila melanogaster* have shown that after 60 generations of adaptation to hot temperatures 75% of genes evolved higher plasticity (Mallard *et al*., 2020), suggesting that adaptation to novel environments leads to an initial increase in plasticity. Altogether, it is still unclear if plasticity precedes and facilitates migration to novel environments, or, if in contrast, exposure to novel conditions initially fosters increased plasticity (Fig. 1), followed by a progressive loss of plastic potential through *genetic assimilation* (Ehrenreich & Pfennig 2016) towards adaptation. Despite their contrasting nature, both scenarios suggest that plasticity plays a pivotal role during early phases of adaptation.

**Figure 1.**
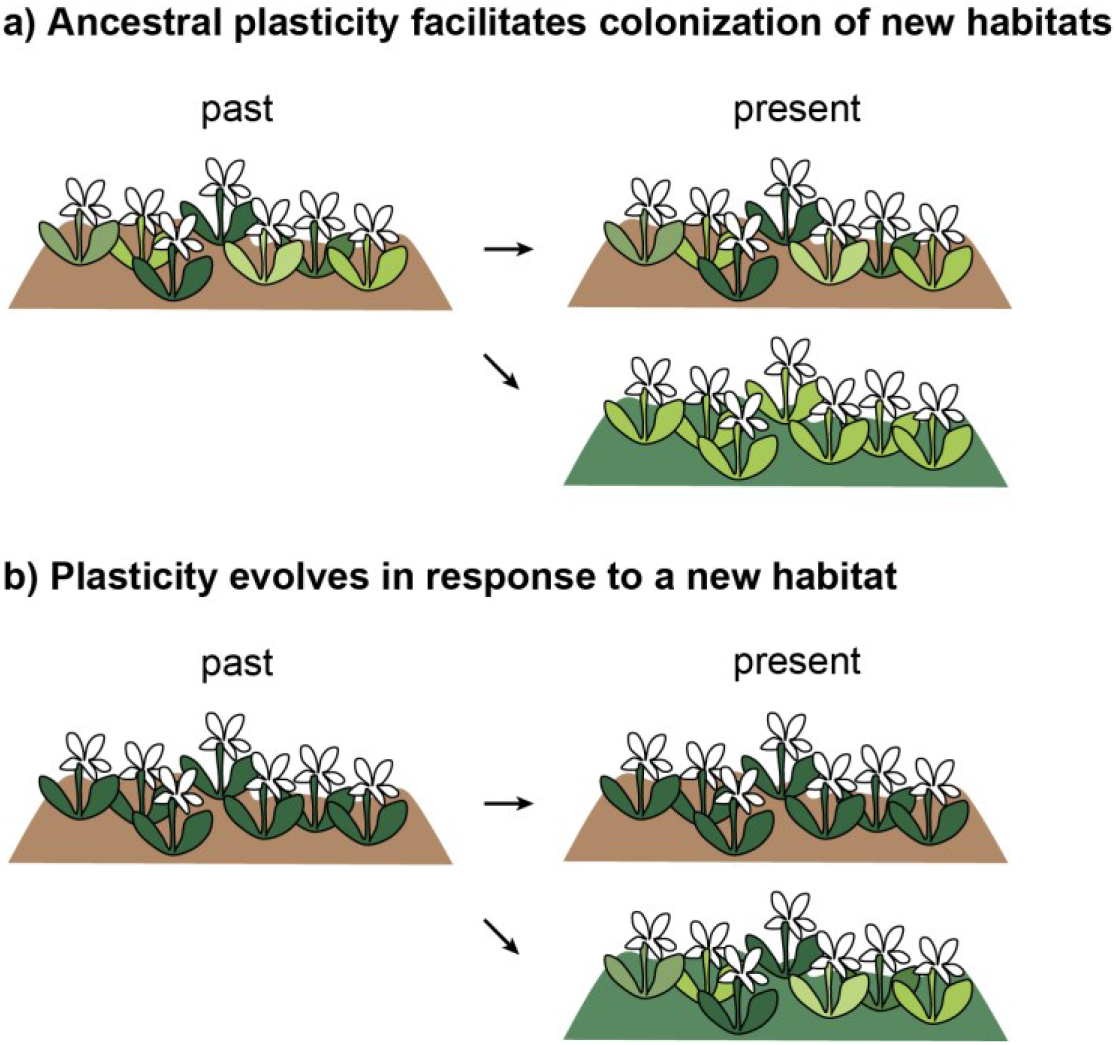
Two hypotheses about the role of phenotypic plasticity during evolution of different ecotypes. The heterogeneity in plant color symbolizes here the degree of plasticity at different stages, whereas light and dark brown indicate an ancestral and a derived niche, respectively. Other aspects, for example the amount of genetic variation in the population, its size, or the temporal and spatial environmental heterogeneity, are not taken into account in these simplified scenarios. **(a)** Pre-existing plasticity in an ancestral population (e.g. in a heterogeneous environment) may facilitate colonization of new habitats. The phenotype is ultimately refined in the newly occupied habitat, where plasticity could be lost over time, due to genetic assimilation. **(b)** The ancestral population bears little plasticity, which evolves in response to a newly colonized environment. This scenario has been coined ‘plasticity-led evolution’ (Schwander & Leimar, 2011; Levis & Pfennig, 2016).

Here, we investigate gene expression plasticity vs. genetically encoded expression differentiation during initial stages of parallel divergence in the plant *Heliosperma pusillum* (Waldst. and Kit.) Rchb. (Caryophyllaceae; Trucchi *et al*., 2017; Szukala *et al*., 2022). This species forms altitudinal ecotypes previously shown to bear cross-generations phenotypic differentiation (Bertel *et al*., 2016b, 2018). The alpine ecotype is widely distributed from the Spanish Cordillera Cantabrica to the Romanian and Ukrainian Carpathians, inhabiting wet screes, rock faces and open grasslands above the timberline, typically at elevations between 1,400 and 2,300 m. In contrast, the montane ecotype (previously referred to as *H. veselskyi* Janka; Frajman and Oxelman 2007; Bertel *et al*., 2016a, 2017, 2018) is restricted to the south-eastern Alps, being represented by isolated and typically small populations, mostly below overhanging cliffs in poor light conditions and shaded from rain in the montane belt (500–1,300 m). Importantly, genomic and transcriptomic studies (Trucchi *et al*., 2017; Szukala *et al*., 2022) have demonstrated, among others with coalescent methods, that pairs of geographically clustered montane and alpine ecotypes in the south-eastern Alps have diverged at least four times independently, representing a case of parallel evolution and can thus be regarded as natural evolutionary replicates.

The lack (alpine) or presence (montane) of a dense indumentum with long multicellular sticky glandular hairs on stem and leaves is the most divergent morphological trait (Fig. 2; Bertel *et al*., 2016a, 2017, 2018). The morphological divergence is most strongly correlated with temperature and soil humidity differences between the two altitudinal sites, whereas humidity and light availability show higher temporal variability at the montane sites compared to the alpine ones (Bertel *et al*., 2018). Along the same lines, montane traits such as multicellular trichomes and physiological response to low light (Bertel *et al*., 2016a) typically show greater variability across and within montane populations (Fig. 2a-c; Bertel *et al*., 2018). This variability is likely at least in part due to plasticity, given that reduced genetic variation was found in the small and strongly confined montane populations (Fig. 2f; Trucchi *et al*., 2017; Szukala *et al*., 2022). Reciprocal transplantation experiments performed at the native altitudinal sites (Bertel *et al*., 2018) demonstrated a home-site fitness advantage of each ecotype in terms of establishment success (i.e. measured as the proportion of plants alive one year after germination). Higher survival rates of either ecotype in its respective native environment are strong indicators that the morphological and physiological differentiation has adaptive value.

**Figure 2.**
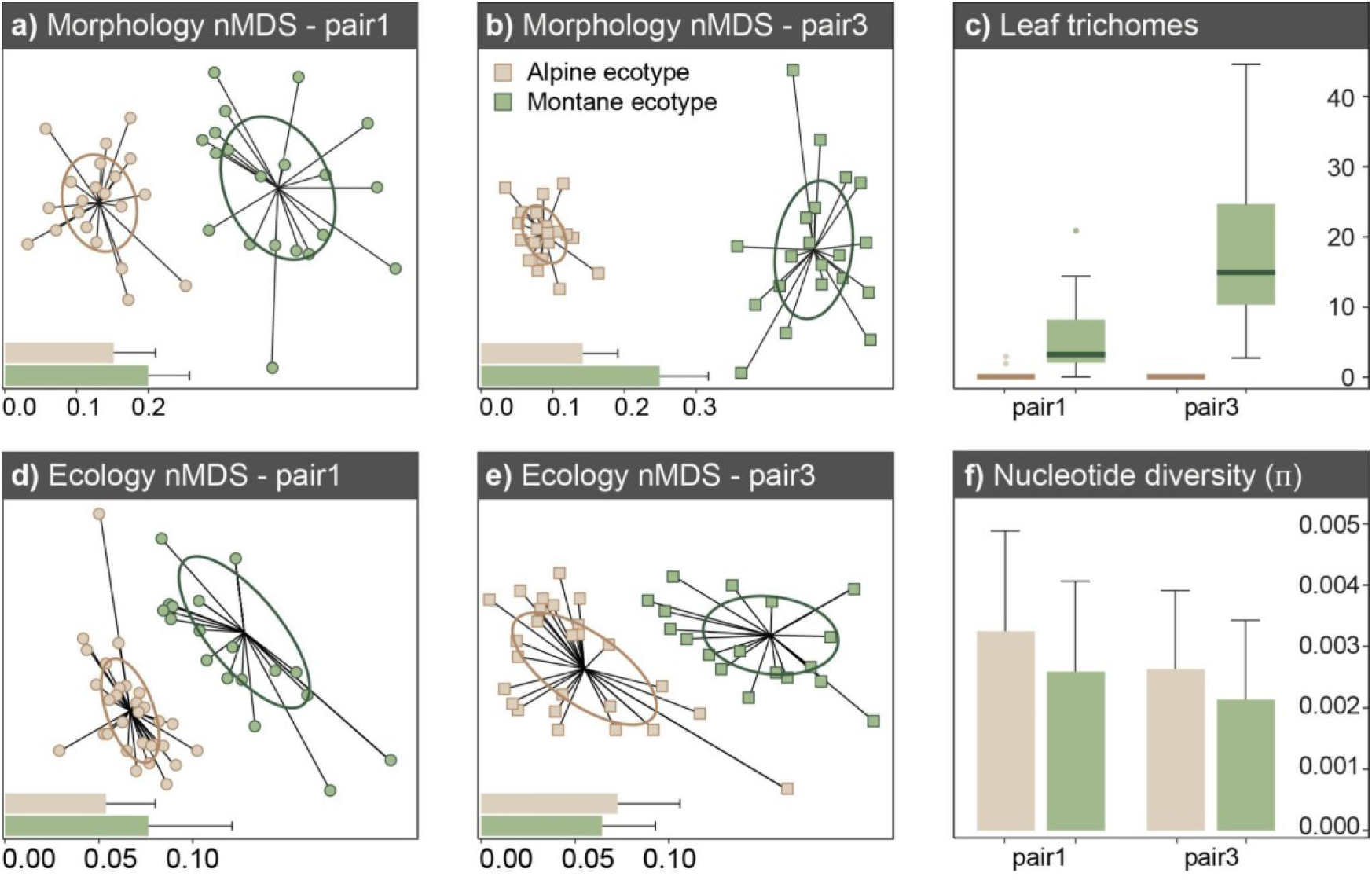
Summary of within-population morphological, ecological and genetic diversity within the two population pairs of *Heliosperma pussilum* investigated here, drawn from previously published data. Green- and brown-filled elements represent the montane and alpine ecotypes, respectively, while circles and squares represent pair 1 and 3, respectively. **(a-b)** Non-metric multidimensional scaling (nMDS) showing increased morphological variability in the montane ecotype among 16 morphometric characters measured on 20 individuals per population in each ecotype and ecotype pair, newly calculated using the data from Bertel *et al*. (2016). Confidence ellipses around treatment centroids represent the standard deviation of the measurements of the respective group. The bar charts on the lower left corner are the mean and SD of the dissimilarity matrices based on Bray-Curtis distances for each population. **(c)** Pronounced variation in the amount of glandular trichomes measured in the montane ecotype, as compared to the alpine, potentially suggestive of increased plasticity in the former. Boxplots drawn from data from Bertel *et al*. (2018). **(d-e)** Non-metric multidimensional scaling (nMDS) of ecological differentiation. Ordinations are based on dissimilarity matrices of mean Landolt indicator values of species growing within a circular area of 0.2 m radius centered on individuals, drawn from data from Bertel *et al*. (2018). **(f)** Estimates of within-population nucleotide diversity (π) in each pair, calculated using RNA-seq data from Szukala *et al*. (2022).

Here, we investigate phenotypic plasticity in the context of initial phases of ecotype divergence in *H. pusillum* (Fig. 1). We quantify gene expression divergence in two parallely evolved ecotype pairs upon reciprocal transplantations at natural growing sites, to identify genes that *i*) diverge in expression between the two ecotypes regardless of the growing environment (*constitutive component of gene expression divergence*), or *ii*) change their expression plastically as a function of the environment (*plastic component of gene expression divergence*), and characterize these two components of ecotype expression differentiation. To reinforce our interpretation of the observed patterns, we investigate the amount of private genetic variation and minor allele frequencies in the ecotypes. Finally, we discuss how plasticity might pave the path towards adaptation during early stages of ecological divergence.

## Results

### Gene expression differences are driven by origin (ecotype pair) and ecotype divergence

To be able to investigate the interaction between altitude and gene expression, we isolated RNA from leaves of two ecotype pairs grown at either the alpine or the montane natural sites (Fig. 3a). The two altitudinal niches are characterized by stark differences in average and amplitude of temperature, water and light availability (Bertel *et al*., 2018), but also by distinct biotic environments (Trucchi *et al*., 2017; Bertel *et al*., 2018). After filtering out genes with low normalized expression counts we searched a total of 15,591 genes for genetically vs. environmentally-driven expression divergence between ecotypes and environments. Multidimensional scaling (MDS, Fig 3b) and principal component analysis (PCA, Supplementary Fig. S1) of normalized read counts showed that gene expression clusters the samples by ecotype pair (i.e., pair 1 shown with circles in Fig. 3b versus pair 3 shown with squares; in Supplementary Fig. S1 PC1 explaining 17.0% of variance), as well as by ecotype (i.e., alpine shown with brown-filled symbols in Fig. 3b versus montane shown with green-filled symbols; in Supplementary Fig. S1 PC2 explaining 10.4% of variance). The variable growing environment explained less variance of the data (montane environment shown with symbols with green margins versus alpine environment shown with symbols with brown margins; in Supplementary Fig. S1 PC5 explaining 4.7% of variance). This clustering shows that the locality of origin of each plant and the divergence between ecotypes explain most of the expression patterns recovered.

**Figure 3.**
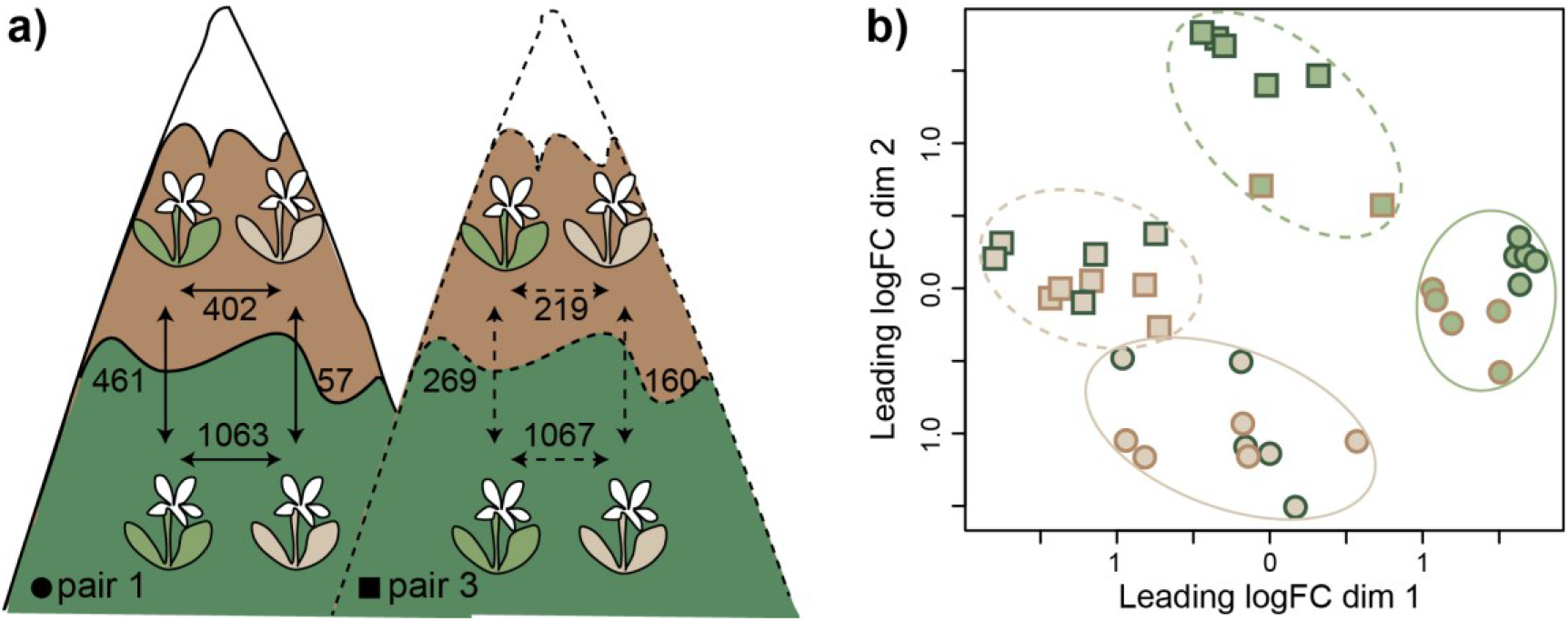
Study design and summary of its results. **(a)** Setup of the reciprocal transplantations between alpine (symbolized with brown areas of the mountains) and montane sites (shown with dark green areas). Brown plants represent the alpine ecotype, green plants the montane one. The numbers represent differentially expressed genes in different comparisons (adjusted p < 0.05). For simplicity the two ecotype pairs are here shown on different mountains, but they have been both reared together at the native localities of pair 1. **(b)** Multidimensional scaling plot of distances between gene expression profiles of individuals of the two ecotypes grown at different altitudes. Circles and continuous lines represent the ecotype pair 1, squares and dashed lines the ecotype pair 3. Green- and brown-filled symbols show the montane and alpine ecotypes, respectively, while green and brown symbol margins represent the low and high growing sites, respectively.

We observed that gene expression differences between ecotypes upon reciprocal transplantation showed similar patterns in both pairs analyzed (Figs. 3–4). In the montane environment, we found 1,063 differentially expressed (DE) genes between the two ecotypes of pair 1 (652 under- and 411 overexpressed in the montane ecotype compared to the alpine one) and 1,067 DE genes between the two ecotypes of pair 3 (483 under- and 584 overexpressed in the montane ecotype compared to the alpine one; green and grey symbols in Fig. 4). By contrast, significantly fewer genes were found to be DE in the alpine environment (brown and grey symbols in Fig. 4) with 402 DE genes between ecotypes in pair 1 (246 under- and 156 overexpressed in the montane ecotype compared to the alpine one) and 219 in pair 3 (83 under- and 136 overexpressed in the montane ecotype compared to the alpine one). Despite these differences in expression patterns observed when comparing the situation between the growing environments, in both ecotype pairs the expression patterns at either altitude were positively correlated (Spearman’s correlation 0.49 and 0.36 in pair 1 and 3, respectively, Fig. 4), implying that an important proportion of gene expression networks is under genetic control and does not change substantially upon environmental change.

**Figure 4.**
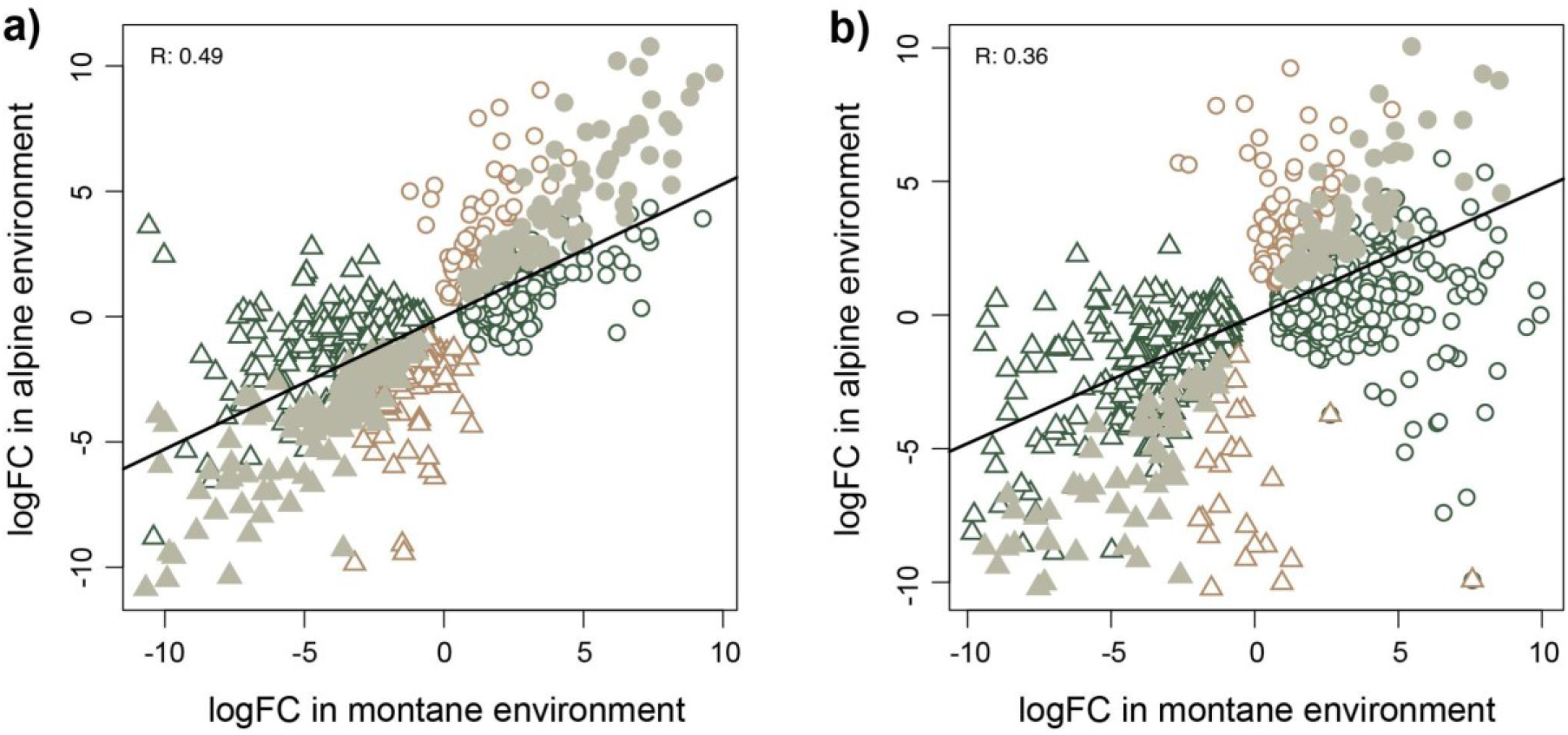
Differentially expressed (DE) genes between ecotypes in different environments. The X- and Y-axis show log fold-change values for genes with differential expression between ecotypes when these are grown in the montane (green and grey symbols) and alpine (brown and grey symbols) environment, respectively, for ecotype pair 1 **(a)** and 3 **(b)**. Triangles and circles represent genes under- and over-expressed in the montane ecotype compared to the alpine, respectively. Grey filled symbols show genes with stable, non-plastic expression divergence between ecotypes, independent of the environment. The black line shows the correlation between gene expression changes in the montane and alpine environments.

### Constitutive evolutionary changes in gene expression

Among the DE genes 216 (more than expected by chance; hypergeometric p < 1e-147) and 118 (more than chance expectations; hypergeometric p < 2.6e-79) genes (grey symbols in Fig. 4) were always DE between ecotypes in the same direction in pair 1 and 3, respectively, regardless of the growing environment. These genes that do not show significant environmental sensitivity represent constitutive expression divergence and are most likely relevant in shaping stable trait differences between ecotypes. Moreover, the genes with constitutive expression divergence appeared to shape a considerable proportion of expression differences in the alpine conditions – i.e., representing consistently ca. 54% of overall expression differences between ecotypes in this environment in both pairs (compared to 20% and, respectively, 11% in the montane environment). Among the constitutive genes identified in each pair, 26 genes (Supplementary Table S1) were shared by both ecotype pairs (more than chance expectations; hypergeometric p < 3e-24). Finally, eight of these genes (Supplementary Table S1) were also found to be DE in both ecotype pairs in a non-native, common garden environment in a previous study (Szukala *et al*., 2022), despite the different growing conditions and developmental stage.

### Environmentally sensitive gene expression

We also looked for environmentally induced expression changes within each ecotype, when these are grown at different elevations (i.e. G×E interaction). We found 461 (pair 1) and 269 (pair 3) DE genes in the montane ecotype versus 57 (pair 1) and 160 (pair 3) DE genes in the alpine ecotype that were explained by the variable “altitude” of the generalized linear model (Figs. 3a and 5). These results suggest that gene expression in the montane ecotype is strongly modified depending on the altitude, implying more pronounced expression plasticity than in the alpine ecotype. This pattern was particularly pronounced in pair 1. In both pairs, the amount of genes showing significant expression plasticity in the montane ecotype is more than double the amount of constitutive DE genes shaping ecotype differentiation. We did not observe a clear pattern of down- or up-regulation of gene expression in the non-native environment that was consistent across both pairs. Also, in both ecotypes, the amount of down-vs. up-regulated genes in the non-native environment was similar.

**Figure 5.**
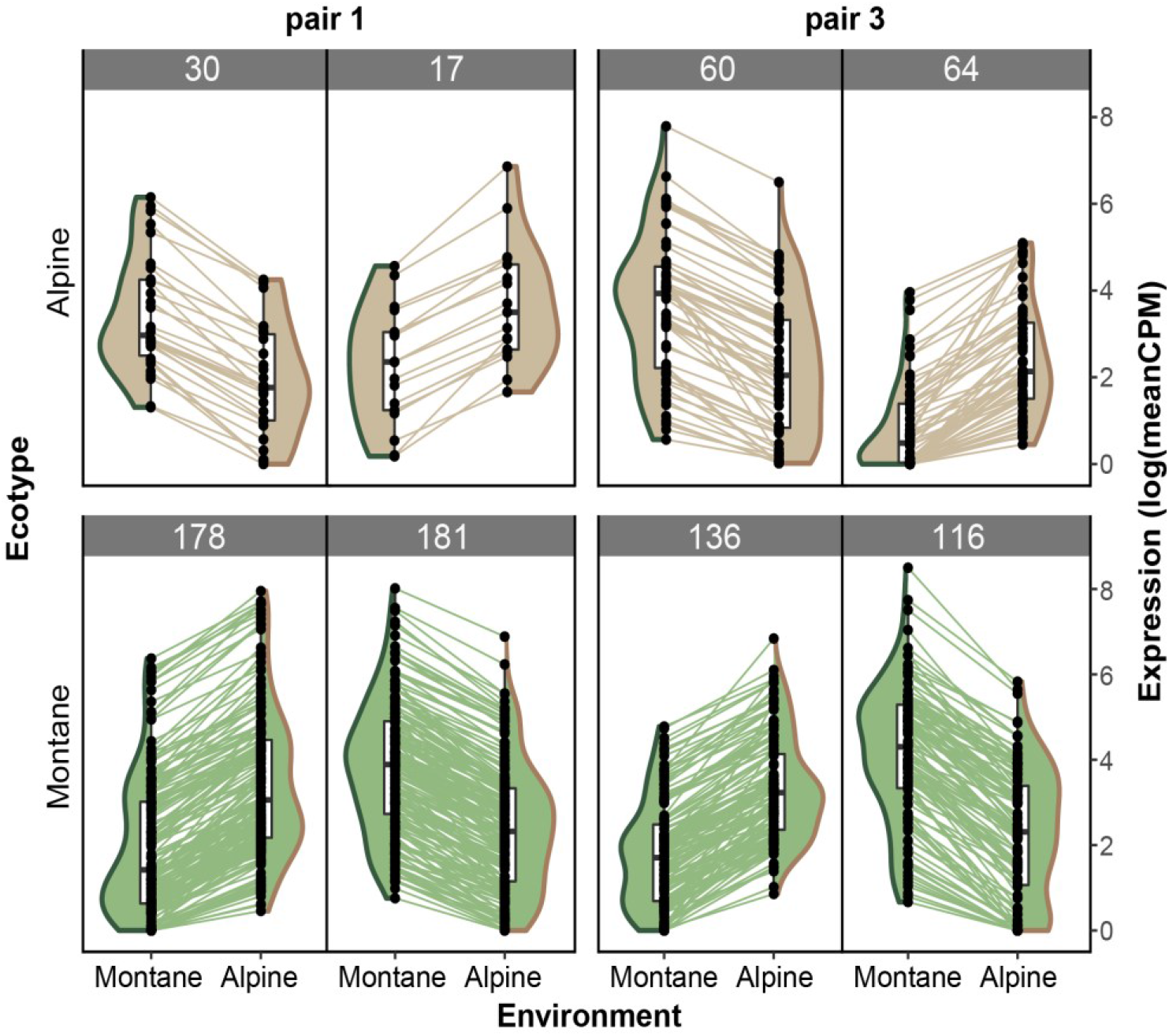
Genotype by environment interactions are more pronounced in the montane ecotype, as exemplified by environmentally driven gene expression changes. Brown- and green-filled violinplots represent the montane and alpine ecotypes, whereas green and brown violinplot margins represent the montane and alpine environment, respectively. Genes DE (adjusted p < 0.05, logFC > 1.5) when alpine (upper row) or montane (lower row) ecotypes are grown at different altitudes are reported. The numbers on top of each plot give the number of genes in the respective category. Each dot represents the average expression of a gene in a given environment, while lines connecting two dots show the expression change of a particular gene at different altitudes.

### Biological significance of constitutive DE genes

The 26 genes consistently DE between ecotypes regardless of the environment and shared by both ecotype pairs are reported with GO term annotations in Supplementary Table S1. The genes overexpressed in the montane ecotype (positive logFC in both pairs) are involved in response to salt-stress and water deprivation (*BSK11, CER1*), epigenetic regulation of gene expression by methylation (*DNMT2*), and protein phosphorylation (*LRR-RLK, RPS20B*), while underexpressed genes in the montane ecotype (negative logFC in both pairs) play roles in immunity (*FUC1, At4g35733, CYP83B1*) and enhanced drought and salt tolerance (*CLB*, see de Silva *et al*., 2011 for increased salt tolerance in knock-out mutants of *Arabidopsis thaliana*).

Additionally, we performed GO terms enrichment of the genes that were consistently DE between ecotypes regardless of the environment (i.e. *constitutive DE genes*) but specific for each ecotype pair separately, to clarify if different sets of constitutive genes do underlie similar functional networks and adaptive responses (Fig. 6a-b, Supplementary Tables S2 and S3). In both pairs we found significant enrichment (adjusted p < 0.05, Fisher-exact test) of response to water deficit and salinity, as well as responses to abscisic acid (ABA), probably related to stress responses (Fig. 6a-b). Despite the convergence of the enriched GO terms, the number of genes underlying each term, as well as the z-score exemplifying the overall expression direction change differed in the two evolutionary replicates (Wolfe *et al*., 2021). Ecologically relevant enriched functions that were not shared by the two pairs have also been identified. In pair 1, DE genes overexpressed in the montane ecotype were enriched for root hair elongation (i.e., a pathway representative also for multicellular trichome development in plants) and epidermal cell differentiation, as well as stomatal closure and negative regulation of gene expression (Fig. 6a). In pair 3, genes overexpressed in the montane ecotype were involved in negative regulation of defense responses, as well as jasmonic acid-mediated signaling (Fig. 6b). The full lists of GO terms enriched in pair 1 and, respectively, pair 3 for genes constitutively DE between ecotypes are reported in the Supplementary Tables S2-S3.

**Figure 6.**
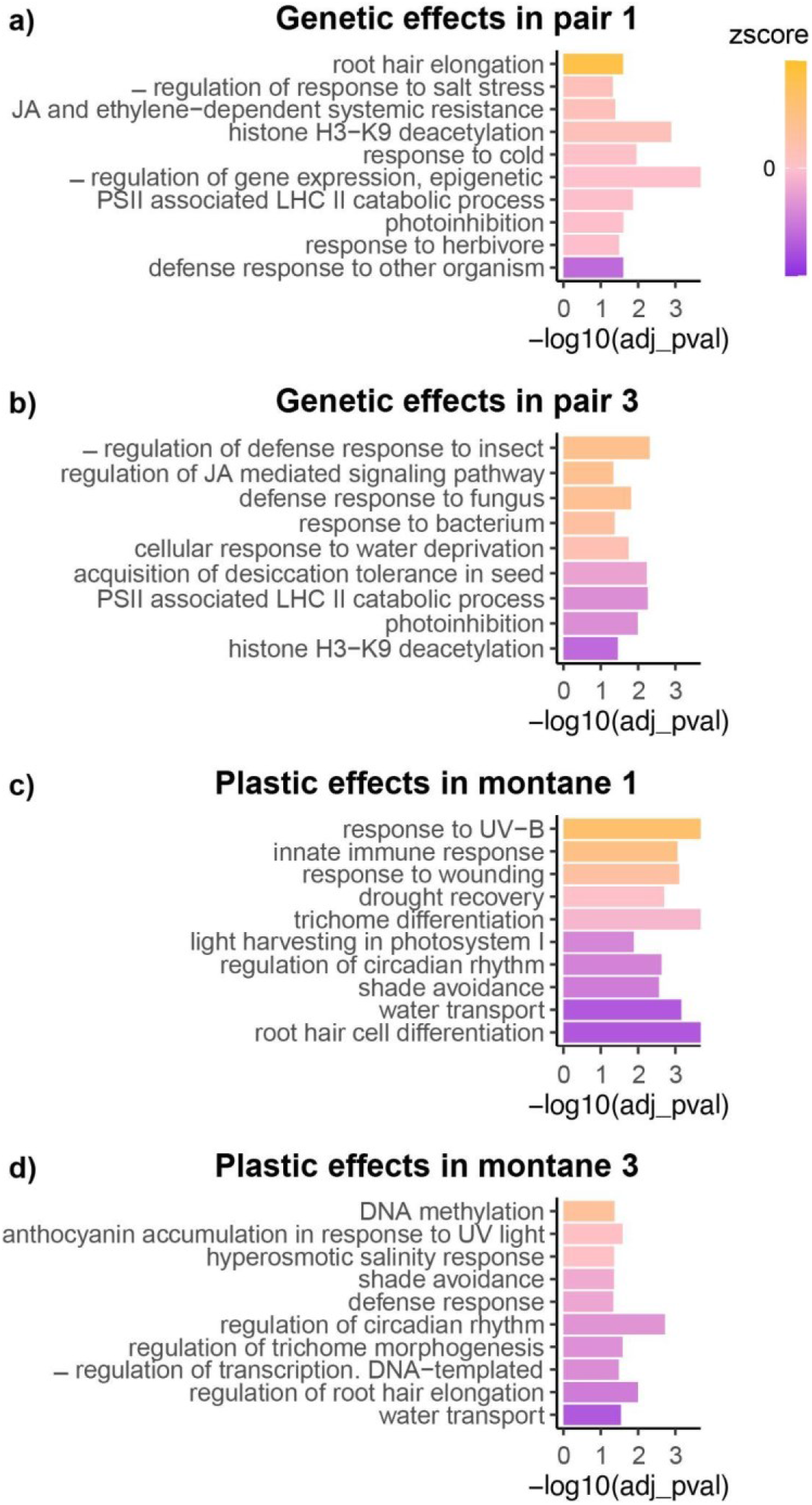
GO terms enrichment (biological processes) of constitutive vs. plastic DE genes. Enriched functions in constitutive expression differentiation between ecotypes in pair 1 **(a)** and 3 **(b)**. Enriched functions in genes changing their expression plastically between altitudes in the montane ecotype 1 **(c)** and 3 **(d)**. Each bar corresponds to a GO term (y axis), while the size of the bars corresponds to the significance of the enrichment (adjusted p < 0.05). The color scale represents the z-score, which is computed based on the logFC of expression of each gene underlying a specific GO term with orange shades corresponding to overexpression in the montane ecotype **(a-b)** or the montane environment **(c-d)**, and violet shades indicate an underexpression in the montane ecotype **(a-b)** or the montane environment **(c-d)**. GO terms reported were selected because of their ecological relevance from a larger list of significant GO terms, fully reported in the Supplementary Tables 2-5. JA, jasmonic acid; LHC, light-harvesting complex; PSII, photosystem II.

### Biological significance of plastic DE genes

We report in Fig. 6c-d, and Supplementary Tables S4 and S5 the GO terms enriched for the genes changing expression plastically in the montane ecotype of both pairs, and in Supplementary Tables S6 and S7 the GO enrichment of the genes changing expression plastically in the alpine ecotype. Terms enriched in plastic DE genes in the montane ecotype were similar among the two ecotype pairs, including response to high light intensity (such as photosynthesis, anthocyanin biosynthesis, response to UV-B and shade avoidance), trichome or root hair differentiation, regulation of circadian rhythm, response to salinity and water transport, and regulation of transcription and methylation. Plastic differential gene expression in the montane ecotype tended to be characterized by pronounced downregulation in the alpine environment in both pairs (Fig. 6c). The direction of expression changes underlying the same function were often inconsistent between pairs, likely depending on the function of different genes affecting the same pathway (Fig. 6c-d). As mentioned above, the alpine ecotype showed reduced plasticity of gene expression compared to the montane one, especially in pair 1. Despite the lower number of DE genes underlying enriched function, we found some similar functions to be enriched as in the montane ecotype (e.g. regulation of circadian rhythm and response to light).

### Population-wise private alleles

After calling and filtering high quality private single nucleotide polymorphisms (SNPs) in each one of the four studied populations we observed an excess of private polymorphisms in the alpine populations compared to the montane ones (Fig. S2a and S3). Additionally, we observed that minor allele frequencies (MAFs) among private alleles tend to be high (i.e., mean(MAF) > 0.2 in all populations, Fig. S2b), suggesting that a majority of them did not accumulate during recent population expansion. We also observed that MAFs of private alleles tend to be similar between montane and alpine populations, suggesting that private variation is not strongly affected by opposite trends in the evolution of the effective population size between ecotypes (i.e. larger alpine populations vs smaller montane ones). Taken together, these results are consistent with previous molecular results (Frajman & Oxelman, 2007; Trucchi *et al*., 2017; Szukala *et al*., 2022) and suggest that the montane ecotypes are derived from the alpine populations of the species.

## Discussion

To date, a handful of studies have investigated the evolution of plasticity during early stages of adaptation in natural (Scoville & Pfrender, 2010; Passow *et al*., 2017; Corl *et al*., 2018; Levis *et al*., 2018) or experimental populations (Huang & Agrawal, 2016; Sikkink *et al*., 2019; Mallard *et al*., 2020; Brennan *et al*., 2021). Here, we have investigated both constitutive and plastic changes in gene expression of altitudinally segregated ecotypes upon reciprocal transplantations in their natural growing sites.

Our results suggest that a combination of constitutive expression divergence and different degrees of expression plasticity underlying the same ecologically relevant functions plays an important role in shaping ecotype divergence. More specifically, the alpine ecotype is characterized by reduced expression plasticity. This ecotype is possibly closer to the ancestral genotype state according to the higher amount of private variation detected in our analyses, which is generally used as a proxy for ancestral state (Schönswetter & Tribsch, 2005; Paun *et al*., 2008). It also experiences extreme environmental factors (e.g. higher amplitude of seasonal temperature fluctuations, low temperatures and enhanced solar irradiation), consistent with the hypothesis that extremes lead to enhanced robustness of gene expression and lack of plasticity (Lande, 2009; Chevin & Hoffmann, 2017). The inability of the alpine ecotype to react plastically to the altitudinal transplantation is possibly consistent with a (albeit not significantly) lower establishment success of alpine plants transplanted to the montane environment compared to transplanted plants of montane origin at alpine elevation (Bertel *et al*., 2018).

By contrast, the derived montane ecotypes bear higher plastic potential of gene expression. As a consequence of this enhanced plasticity, the expression profiles of the ecotypes were more similar in the alpine environment, while they differed strongly in the montane one, except for a minor proportion of constitutive expression changes. Similar to our results, enhanced phenotypic plasticity in low-elevation individuals compared to high-elevation ones was found in *Wahlenbergia ceracea* by Nicotra *et al*. (2015). In this species, higher plasticity in low-elevation plants was shown to be adaptive, whereas plasticity in high-elevation plants was more likely to be maladaptive. In the same study, higher epigenetic diversity in response to growth temperature detected in seedlings from low elevation suggested a role for DNA methylation in shaping adaptive plastic responses. Possibly in line with these results, our GO terms enrichments (Fig. 6, Supplementary Tables S2-6) showed that both constitutive gene expression divergence, as well as gene expression plasticity are enriched for epigenetic processes (e.g. histone H3-K9 deacetylation and methylation in constitutive and plastic DE genes, respectively). However, a previous proof-of-concept study did not reveal significantly different within-ecotype levels of genome-wide DNA methylation variation between the two ecotypes (Trucchi *et al*., 2016).

Similarly as many alpine plant species (Giesecke *et al*., 2017), *H. pusillum* likely migrated upslope after the Last Glacial Maximum (LGM) likely tracking their specific alpine conditions; the few montane populations present today could represent relicts of the original LGM populations. The alpine ecotype was resolved as ancestral in the study of Frajman and Oxelman (2007) and the time of divergence between ecotypes was estimated around LGM (Trucchi *et al*., 2017), albeit older in another study (Szukala *et al*., 2022). The relict montane populations likely adapted to the specific niche under overhanging cliffs with lack of competition but high levels of abiotic stress (Davis, 1951; García & Zamora, 2003, Minuto *et al*., 2012), as this progressively became warmer and the original alpine habitats at low elevations strongly reduced due to the advancement of forests during the Holocene. The alpine ecotype, on the other hand, enlarged its distribution range throughout the southern European mountain ranges, where its habitats are abundant. Despite both habitats likely differing from the ancestral environment preceding ecotype divergence, we hypothesize that the alpine niche resembles more the ancestral one with regard to temperature and humidity, as well as biotic interactions. Under this scenario, the montane ecotype have enhanced expression plasticity as a consequence of exposure to a new niche in the montane environment, starting from an ancestral state, in which plasticity was lacking. Alternatively, the alpine ecotype might have lost ancestral plasticity through canalization. Although this second hypothesis appears less likely, it cannot be safely ruled out.

Despite some differences in the magnitude of the patterns found, our results were consistent between the two ecotype pairs analyzed, which represent independent instances of ecotype formation (Trucchi *et al*., 2017; Szukala *et al*., 2022), and can therefore be considered natural evolutionary replicates. It is important to notice that we could sample only two biological replicates of the montane ecotype from pair 3 transplanted to the alpine site. Interpretations of the results regarding this group should therefore be considered with caution. Still, while differences in the absolute numbers observed between ecotype pairs might have been driven by these differences in sampling density, the overall patterns of evolutionary vs. plastic expression changes should not be affected severely.

Since evolution favored the maintenance of enhanced plasticity in the montane environment in at least two independent divergence events, we hypothesize that enhanced plasticity leads to a fitness advantage in the incipient stages of adaptation to new habitats (i.e., in this system, the montane habitats; Bertel *et al*., 2018). Plasticity might indeed be beneficial in the stressful montane niche under overhanging cliffs which is characterized by heterogeneous light and water availability, longer periods of drought, and higher average temperatures (Bertel *et al*., 2018). Our results align with experimental studies showing that adaptation to novel conditions (e.g. high temperature) increases gene expression plasticity (Mallard *et al*., 2020; Brennan *et al*., 2021). Moreover, exposure to abiotic stress, such as drought, salinity, and heat, induced high gene expression plasticity in *Brachypodium distachyon* (Priest *et al*., 2014). Like in *Heliosperma*, convergence in the evolution of plasticity was found in two parallelly evolved zinc-tolerant lineages of *Silene uniflora* (Wood *et al*., 2021). Nevertheless, zinc-tolerant *Silene* derived populations appeared to have decreased plasticity due to genetic assimilation of ancestral plasticity. Future studies should aim to directly assess if expression plasticity in the montane ecotype changes the phenotype deterministically in such a way that fitness is increased, in order to drive stronger conclusions about the impact of natural selection on plasticity in this system. Understanding the importance of phenotypic plasticity for fast adaptation to abiotic stress is very timely also for crops and breeding (Shao *et al*., 2007; Dalal *et al*., 2017; Fox *et al*., 2019).

We found that over 50% of the genes DE between ecotypes in the alpine environment were also DE in the montane environment, implying that an important part of expression divergence in the alpine environment is driven by evolutionary change, while a major additional proportion of divergence in the montane environment is plastic. Consistent with a previous investigation of ecotype-specific gene expression profiles in a common garden (Szukala *et al*., 2022), we found a limited, but still significant amount of constitutive DE genes shared by the two ecotype pairs. This result confirms the previously observed heterogeneity of DE genes in parallely evolved ecotype pairs, suggesting redundant adaptive solutions to cope with altitudinal differentiation. Interestingly, eight among these 26 genes were previously found to be DE in both pairs in Szukala *et al*. (2022), even if the seeds were collected in a different year and RNA was extracted from leaves at a different developmental stage and in different environments. Notably, we recovered similar biological functions enriched for constitutive DE genes, especially related to (a-)biotic defense responses, such as herbivory, temperature, water deprivation and salt-stress (note that the last two stressors are functionally strongly interconnected; Ma *et al*., 2020), light availability, and epigenetic regulation, despite the limited overlap of specific constitutive genes. Similar functions related to the morphological (i.e. differences in hairiness) and ecological (i.e. differences in temperature, and water and light availability) divergence of the populations were underlied by both plastic and constitutive gene expression divergence.

In summary, the comparison of gene expression patterns between ecotypes upon reciprocal transplantations provided insights into the relative roles of expression plasticity and evolution in shaping gene expression divergence in nature. Similarly as in precedent studies (McCairns & Bernatchez, 2010; Narum & Campbell, 2015), our findings point to an intricate interaction of evolutionary changes and plasticity, and to an important role of expression plasticity favoring the colonization of novel habitats during early stages of divergence. Future studies should aim for a better understanding of the regulatory patterns behind plasticity and its role in shaping adaptation.

## Materials and Methods

### Reciprocal transplantations and plant material

Reciprocal transplantations were carried out in 2017 in Lienzer Dolomiten, Kärnten (Austria; alpine site: 46.762 N 12.877 E, 2,055 m; montane site: 46.774 N 12.901 E, 790 m). Seeds were collected from wild populations of both ecotypes at these two localities and in the Puez-Geisler region, Trentino-Südtirol/Alto Adige (Italy; alpine site: 46.601 N 11.768 E, 2290 m; montane site: 46.564 N 11.77 E, 1690 m). We use the same acronyms as in Bertel *et al*. (2018) and Szukala *et al*. (2022), and name the ecotype pair from Puez-Geisler as pair 1, and that from Lienzer Dolomiten as pair 3, to facilitate comparisons between studies. Seeds were first germinated in a common garden in the Botanical Garden of the University of Innsbruck, Austria, and young seedlings were then transferred to the transplantation sites and grown for one season before sampling leaves in early autumn 2017. This approach was necessary, as transplantation trials attempting germination directly at the native sites showed insufficient and erratic germination rates, especially in the dry montane habitats (Bertel *et al*., 2018).

Our approach included hundreds of individuals, and originally aimed to finally investigate with RNA-seq a total of 40 individuals: two ecotypes × two pairs × two elevations × five individuals. However, due to the death of individuals during the course of the experiment (i.e., the alpine site experienced pronounced damage by chamois, among others), only two individuals remained available for the group of montane individuals from pair 3 transplanted to the alpine site. For this reason, our final analyses comprise a total of 37 individuals with five biological replicates per group, and one group with only two biological replicates.

### Library preparation and sequencing

Fresh vegetative shoots from transplanted and control plants were fixed in RNAlater (Sigma) on the same day and time of the day (ca. 2:00 pm) and stored at -80 °C until further processing. Total RNA was extracted from *ca* 90 mg leaves using the mirVana miRNA Isolation Kit (Ambion) following the manufacturer’s instructions, and it was further depleted of residual DNA with a RNase-Free DNase Set (Qiagen) and of the abundant ribosomal RNA by using a Ribo-Zero rRNA Removal Kit (Illumina). RNA was quantified with a NanoDrop2000 spectrophotometer (Thermo Scientific), and its quality assessed using a 2100 Bioanalyzer (Agilent). NEBNext Ultra Directional RNA Library Prep Kit (New England Biolabs) was used to prepare strand-specific libraries. Individually-indexed libraries were pooled together and sequenced with single-end reads (100 bp) on two runs of Illumina NovaSeq S1 at the Vienna Biocenter Core Facilities (VBCF; https://www.viennabiocenter.org/facilities/).

### Differential expression analyses

After demultiplexing using BamIndexDecoder v.1.03 (available from http://wtsi-npg.github.io/illumina2bam/#BamIndexDecoder), bam files were converted to fastq using samtools v.1.3 (Li *et al*., 2009) and quality and adapter trimmed using trimmomatic v.0.36 (Bolger *et al*., 2014). The individual samples were aligned against the reference genome for *Heliosperma pusillum* v.1.0 (Szukala *et al*., 2022) using the available .gff file for gene annotations and STAR v.2.6.0c (Dobin *et al*., 2013). A table of counts was produced using FeatureCounts v.2.0.3 from Rsubread package (Liao *et al*., 2014) including only uniquely mapping reads. After filtering count matrices retaining genes with an average count per million higher than 1, data normalization and differential expression (DE) analyses were performed using the Bioconductor package EdgeR v.3.24.3 (Robinson *et al*., 2010) implementing a generalized linear model of the type *expression = pair + altitude + ecotype + pair*altitude*ecotype* to account for the effects of the covariates altitude, ecotype pair and ecotype on gene expression. Gene-wise dispersion was estimated over all genes using the *estimateDisp()* function and specifying *robust=T* to robustify the estimation against potential outliers. We fitted a gene-wise negative binomial generalized log-linear model (EdgeR function *glmFit*), again with the option *robust=T* to decrease the informativeness of outlier genes. A likelihood ratio test (EdgeR function *glmLRT*) was used to test for DE genes and the significance was adjusted using Benjamini-Hochberg correction of p-values to account for multiple testing. Spearman correlation tests between gene expression changes at different altitudes were also performed.

First, we looked for DE genes between ecotypes in one environment and across both environments (i.e., *constitutive expression divergence*). Second, we aimed to detect plastic expression changes due to the component altitude in each of the ecotypes in each pair. We checked which genes are DE between altitudes in each ecotype, and additionally identified the genes showing a minimum mean fold change (FC) in expression of 1.5 across biological replicates when the growing environment is changed. The FC threshold was set to detect genes showing a strong association with the environmental change. Finally, we tested the statistical significance of the overlap between lists of DE genes using the genes retained after trimming low counts as background and the hypergeometric test of the Bioconductor package SuperExactTest (Wang *et al*., 2015).

### Biological interpretation of DE genes

To retrieve functional annotations of the genes, we updated the functional annotations of the gene models for the reference genome v.1.0 for *Heliosperma pusillum* (Szukala *et al*., 2022) by blasting against the latest *Arabidopsis thaliana* database using Blast2GO v.5.2.5 (Götz *et al*., 2008). Fisher’s exact tests implemented in the Bioconductor package topGO v.2.34.0 (https://bioconductor.org/packages/release/bioc/html/topGO.html) was used to identify significantly overrepresented functions (adjusted p < 0.05).

### Detection of population-wise private alleles

We sorted mapped files according to the mapping position, and marked and removed duplicates using Picard v.2.9.2 (https://broadinstitute.github.io/picard/). Variant calling was then performed following standard practices for RNA as implemented in GATK v.4.1.8.1 (Van der Auwera & O’Connor, 2020). First, reads with Ns in the CIGAR string were split using the split’N’trim function and overhangs were trimmed. HaplotypeCaller was used to call variants with the option *-ERC GVCF*. Subsequently, multiple samples in gvcf format were merged using the GenomicsDBImport utility with the *-L* option to operate in parallel on multiple genomic intervals. Finally, we used GenotypeGVCFs to perform joint genotyping. We filtered the obtained vcf file first using the vcfallelicprimitives modality implemented in vcflib v.1.0.2 (https://github.com/vcflib/vcflib) with the options *--keep-info --keep-geno* to split multiple nucleotide polymorphisms (MNPs) into multiple SNPs. VCFtools v.0.1.16 (Danecek *et al*., 2011) was used to keep only high-quality biallelic SNPs with the options -*-max-alleles* 2 *--min-alleles* 2 *--minDP* 4 *--minGQ* 20 *--minQ* 30 *--remove-indels*. Additionally, we filtered using the *--max-missing* 1 option to discard all loci with missing genotypes and test for consistency of the results when missingness was not allowed. To detect population-wise private alleles we used the vcf-contrast module of VCFtools with the options *-n -f -d* 5 and specifying the population samples using *--indv*. Raw numbers of private alleles per population were normalized by the number of samples in each ecotype and population (i.e., ten individuals, except for the montane ecotype of pair 3 with seven individuals). Finally, VCFtools was run on the output files of vcf-contrast with the option *--freq* and specifying the population samples using *--indv* to obtain major and minor allele frequencies for the private alleles.

## Acknowledgments

This work was financially supported by the Austrian Science Fund (FWF) through the doctoral programme (DK) grant W1225-B20 to a faculty team including O.P., and through grant Y661-B16 to O.P. We thank Nicholas Barton, Andrew Clark, Virginie Courtier-Orgogozo, Joachim Hermisson, Thibault Leroy, Magnus Nordborg, John Parsch, Christian Schlötterer for insightful comments and feedback. We thank Marie Huber and Daniela Paun for their support during laboratory work and data acquisition, Pau Cornicero Campmany for help during setting up of the reciprocal transplantations, as well as Martina Imhiavan and Daniel Schlorhaufer from the Botanical Garden of the University of Innsbruck for germinating the plants. Computational resources were provided by the Vienna Scientific Cluster (VSC) and the Life Science Compute Cluster (LiSC) of the University of Vienna. A permit to conduct the presented research activities was granted by the Parco Naturale Dolomiti Friulane (no. 1943); for the Austrian federal state Tirol no such permit was necessary.

## Author Contributions

Study conceived and designed by OP, BF and PS. Sampling of seeds was performed by BF and PS. Transplantation experiments were performed by OP and PS. Bioinformatics and statistical analyses were conducted by AS, with contribution from CB for vizualization of Fig. 2. The manuscript was drafted by AS, and was revised and approved by all authors.

