## Supplementary Fig. for "Parallel adaptation to lower altitudes is associated with enhanced plasticity in *Heliosperma pusillum* (Caryophyllaceae)"

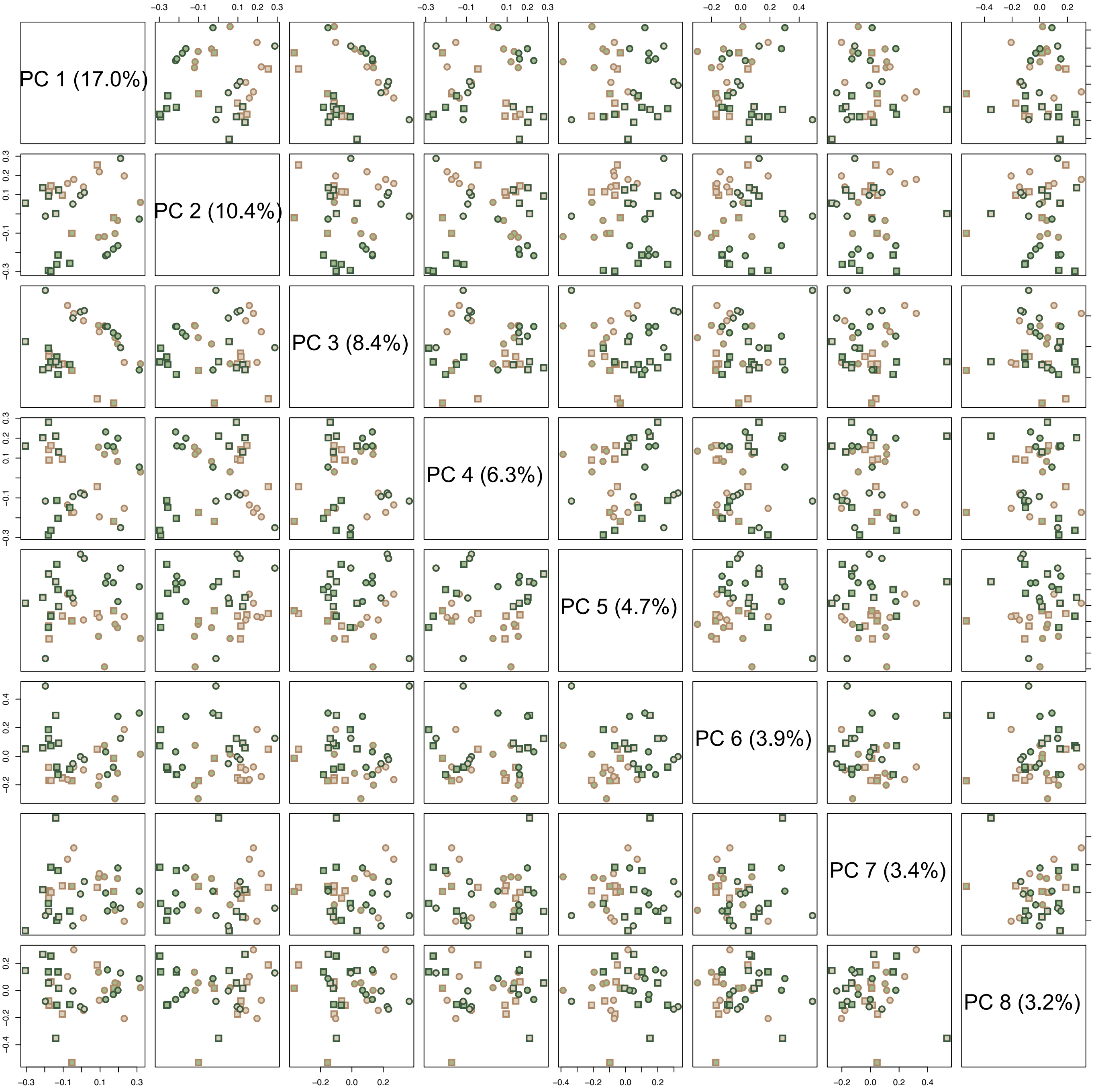


**Supplementary Figure S1. Main trends in normalized gene expression counts.** Principal component analysis of normalized gene expression counts. Visualization of the components 1 to 8. Circles represent the ecotype pair 1, squares the ecotype pair 3. Green- and brown-filled symbols show the montane and alpine ecotypes, respectively, while green and brown symbol margins represent the low and high growing sites, respectively.

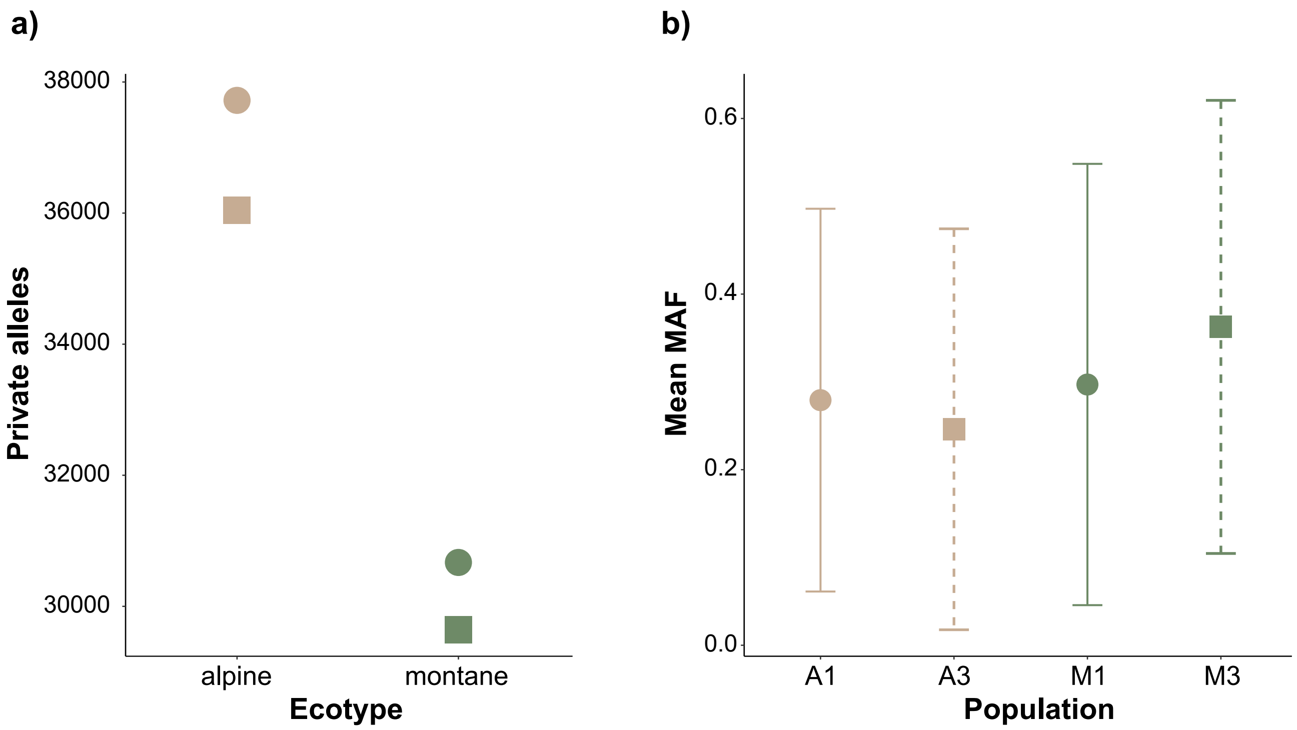


**Supplementary Figure S2. Population private alleles and their minor allele frequency (MAF).** **(a)** Total amount of private alleles by population after normalization by sample size and considering only biallelic SNPs (no missing data allowed). **(b)** MAF of private alleles by population. Bars represent the standard deviation.

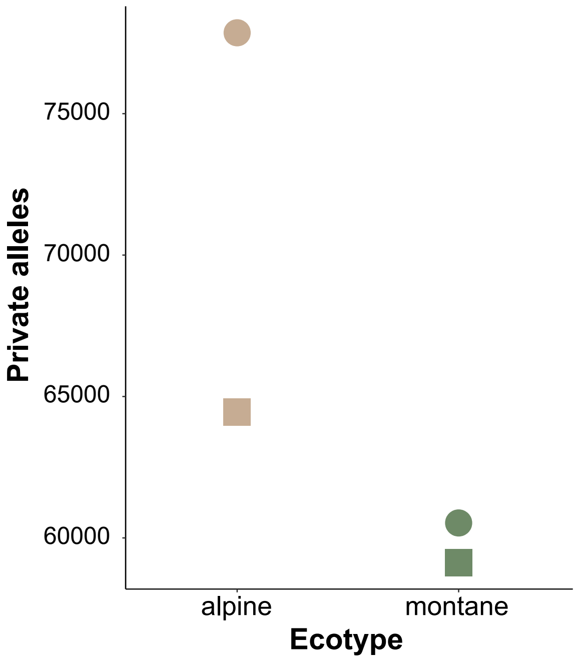


**Supplementary Figure S3. Per population private allele statistics when allowing missing data.**
